# Sysrecon: A systematic data-driven tool for genome-scale metabolic reconstructions

**DOI:** 10.1101/2023.12.13.571370

**Authors:** Shilin Ouyang, Zihao Li, Jiamin Hu, Miyuan Cao, Xiting Wang, Hongxia Sun, Feng Yu, Longfei Mao

## Abstract

Genome-scale metabolic reconstructions contain the information about metabolism, which understands organisms’ molecular mechanisms better. Different reconstruction procedures are developed to achieve high-quality GSMRs of organisms; however, the descriptions of procedures are vague, meaning that they cannot be qualitatively reproduced, even contradictory, and different published reconstructions may involve different steps. This indicates that there is no unified strategy to quantify the steps in GSMRs. To resolve the problem, we created a R package, Sysrecon, to quantitatively and qualitatively describe the steps of GSMRs in the literature. Sysrecon disassembles GSMR into 93 steps, which are compiled after the comprehensive textual analysis of metabolic reconstructions. Then, Sysrecon decomposes each step of the reconstruction procedure into three components: step content, step conversion, and database and tools. When building a GSMR, Sysrecon creates a template procedure that includes a list of pipeline steps, the description of the information conversion in the steps, and the database and tools that may be used to facilitate the conversion. Because each step’s conversion is defined by a formula, the entire GSMR can be dismantled into many ‘step’ blocks. The reconstruction procedure is therefore transformed into an adaptable automated assembly line that can be customized and tailored to settings of different organisms. To the best of our knowledge, Sysrecon is the first computational tool that dynamically constructs the GSMRs pipeline and provides a framework for defining reconstruction steps and can be used as a basis for the development of high-quality GSMRs.

## Introduction

A genome-scale metabolic reconstruction (GSMR) is a comprehensive and systematic representation of the metabolic processes that occur within an organism, based on its genome sequence. This reconstruction integrates information from various biological databases, biochemical literature, and experimental data to create a network of biochemical reactions and associated genes and enzymes (Edwards and Palsson, 1999). Since the first GSMR of *Haemophilus influenzae* RD was reported in 1999, GSMR has been established as one of the major modeling approaches for systems-level metabolic studies (Edwards and Palsson, 1999). GSMRs now include an increasing number of species, not only single-cell species, such as *Escherichia coli* (Edwards and Palsson, 2000; Monk et al., 2017; Orth et al., 2011) and *Saccharomyces cerevisiae* (Förster et al., 2003; Nookaew et al., 2011; Hu et al., 2023), but now multicellular organisms, such as humans(Duarte et al., 2007; Thiele et al., 2013; Brunk et al., 2018; Robinson et al., 2020; Wagner et al., 2021) and plants (de Oliveira Dal’Molin et al., 2010; Mintz-Oron et al., 2012; Pfau et al., 2018a; Moreira et al., 2019; Saha et al., 2011; Shaw and Cheung, 2018; Simons et al., 2014; Gerlin et al., 2021), are also reconstructed. GSMRs allow the prediction of metabolic flux values for metabolic reactions using optimization techniques such as Flux Balance Analysis (FBA) with linear programming (Orth et al., 2011) and thus have many applications in scientific research and industrial production, such as prediction of drug targets(Kim et al., 2016; Mishra et al., 2018; Guo et al., 2022), prediction of enzyme function (Oberhardt et al., 2016) and understanding human diseases (Aller et al., 2018; McGarrity et al., 2018).

A growing number of protocols (Thiele and Palsson, 2010), source tracking and interoperability guidelines (Heavner and Price, 2015; van Dam et al., 2019; Wilkinson et al., 2016), databases (Caspi et al., 2016; Christopher S. Henry et al., 2010; Kanehisa et al., 2012; Satish Kumar et al., 2007b; Schellenberger et al., 2010) and algorithms (DeJongh et al., 2007b; Heirendt et al., 2019a; Christopher S. Henry et al., 2010; Machado et al., 2018b; Satish Kumar et al., 2007b; Wang et al., 2018b) have been developed to facilitate the process of collecting, organizing, and integrating vast amounts of biochemical knowledge. However, building a high-quality GSMR is still time-consuming and prone to missing important metabolic contents. Ambiguous annotations still exist with manual management. For instance, GSMR process commonly suffers from a lack of organic cofactors (Xavier et al., 2017). Similarly, metabolic annotations were missing in roughly 60% of the 99 metabolic reconstruction models (Haraldsdóttir et al., 2014). While these problems can currently be identified and corrected using the functions provided in the COBRA toolbox (Heirendt et al., 2019) or the GlobalFit algorithm (Fritzemeier et al., 2017), a tool that guides and evaluates metabolic reconstruction can minimize these problems.

Tools for standardized metabolic reconstruction quality assessment, such MEMOTE, have recently been created (Lieven et al., 2020). MEMOTE has made some efforts to measure reconstruction quality in four areas: component namespaces, biochemical consistency, network topology and version control. MEMOTE, however, does not have the ability to investigate and analyze the procedures involved in metabolic reconstruction for evaluation and it solely concentrates on the grammatical accuracy of the metabolic reconstruction. More importantly, the steps used to describe the metabolic reconstruction of different species vary widely throughout the relevant literature. This can be attributed to the fact that the metabolic reconstruction steps, data transformation and tools required vary for each species. Current studies have not investigated the preferences of reconstruction steps, transformation methods, and tools used by different organisms in performing GSMR, making it difficult to provide a solution for reconstructing a high-quality GSMR from the three levels: reconstruction steps, transformation methods and tools needed.

Here, we developed an R package Systematic genome-scale metabolic reconstruction (called Sysrecon) (**Figure 1**), which is designed to provide metabolic reconstruction steps, transformations, tools for GSMR of different species. Sysrecon can help ones to assist GSMR. We quantitatively analyzed the different GSMRs, obtained the elemental ratio of each variable in the different reconstructions, and quantitatively compared the different reconstructions based on their ratio. Our study provides a systematic framework for reconstructing high-quality GSMR based on the evaluation of metabolic reconstruction steps.

**Figure 1:**
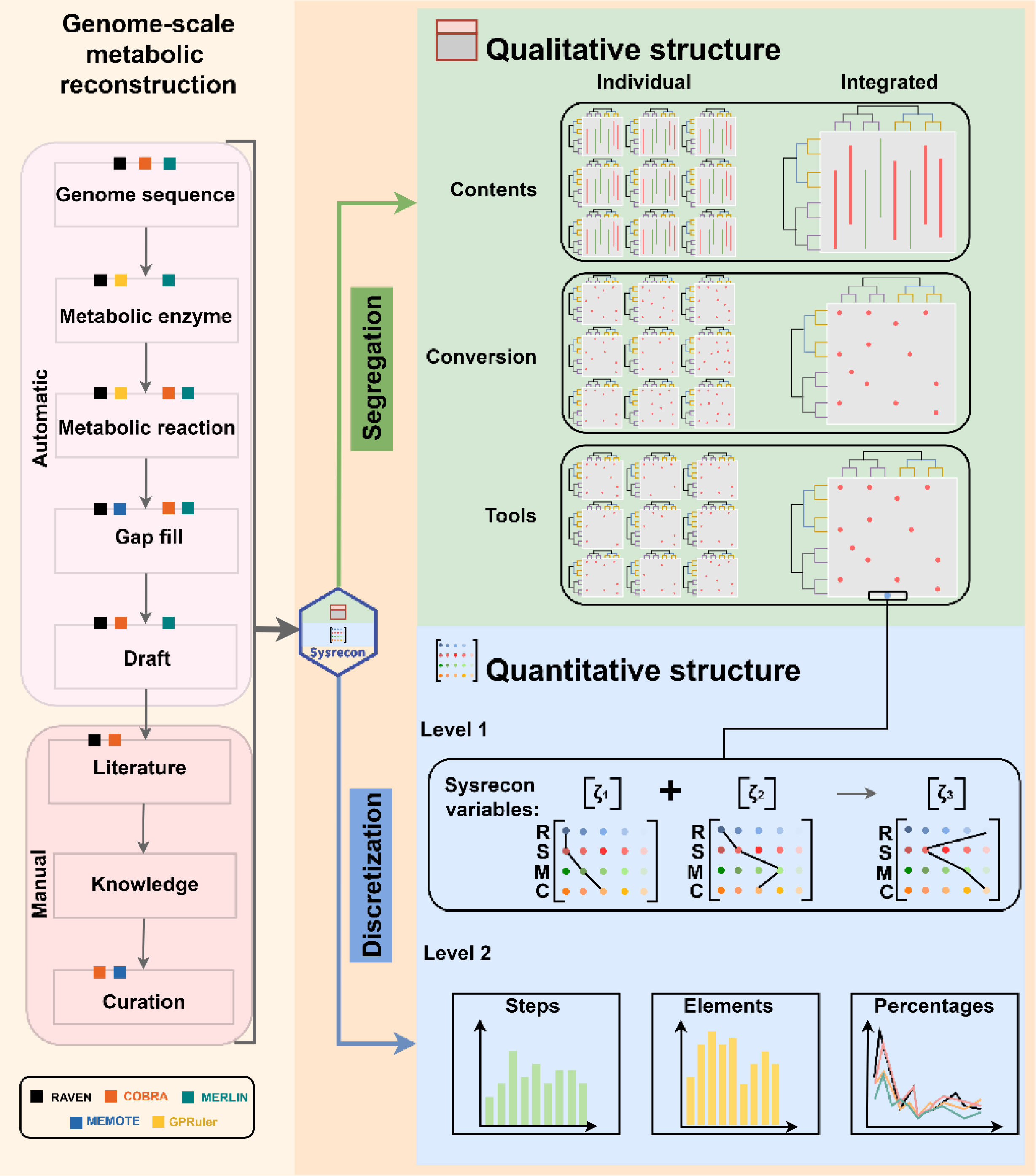
The flow chart of Sysrecon. There are two types of GSMRs: automated generation steps that utilize relevant software or packages to retrieve data, and manual steps that typically require manual data acquisition. Upon inputting the contents of a genome metabolism reconstruction article into the Sysrecon package, qualitative and quantitative results can be obtained. The qualitative findings essentially illustrate the reconstruction procedures, tools, and transformations presented in the essay. In contrast, the quantitative results quantify the reconstruction steps and elements included in the article, as well as perform elemental analysis to identify commonalities and specificities among different GSMRs.

## Methods

### Collecting literatures

We collected 175 literature sources on metabolic reconstruction from Gu et al.’s supplementary materials(Gu et al., 2019) and PubMed, comprising 10 articles on archaea, 110 articles on bacteria, and 55 articles on eukaryotes (**Supplementary data**). These literature sources spanned various publication years (from 2004 to 2018) and employed diverse methodologies, facilitating a comprehensive analysis of metabolic reconstruction across different species. Subsequently, to mitigate the risk of inaccuracies stemming from automated extraction, we manually extracted the content pertaining to metabolic reconstruction from these literature sources.

### Processing text

To count and evaluate the vocabulary in the text, *get_term_matrix* function was developed using the *tm* package, which is based on the text mining (Feinerer et al., 2008). First, the corpus consisted of input texts via *tm’s VCorpus* function with default parameters. Then, the corpus was preprocessed to remove extraneous data by *tm_map*, including the removal of deactivation words and the change of capitalization to lowercase. Finally, using the function *TermDocumentMatrix*, the data in the corpus was converted into a matrix of word item frequencies, namely words matrix (**Figure 2**).

**Figure 2:**
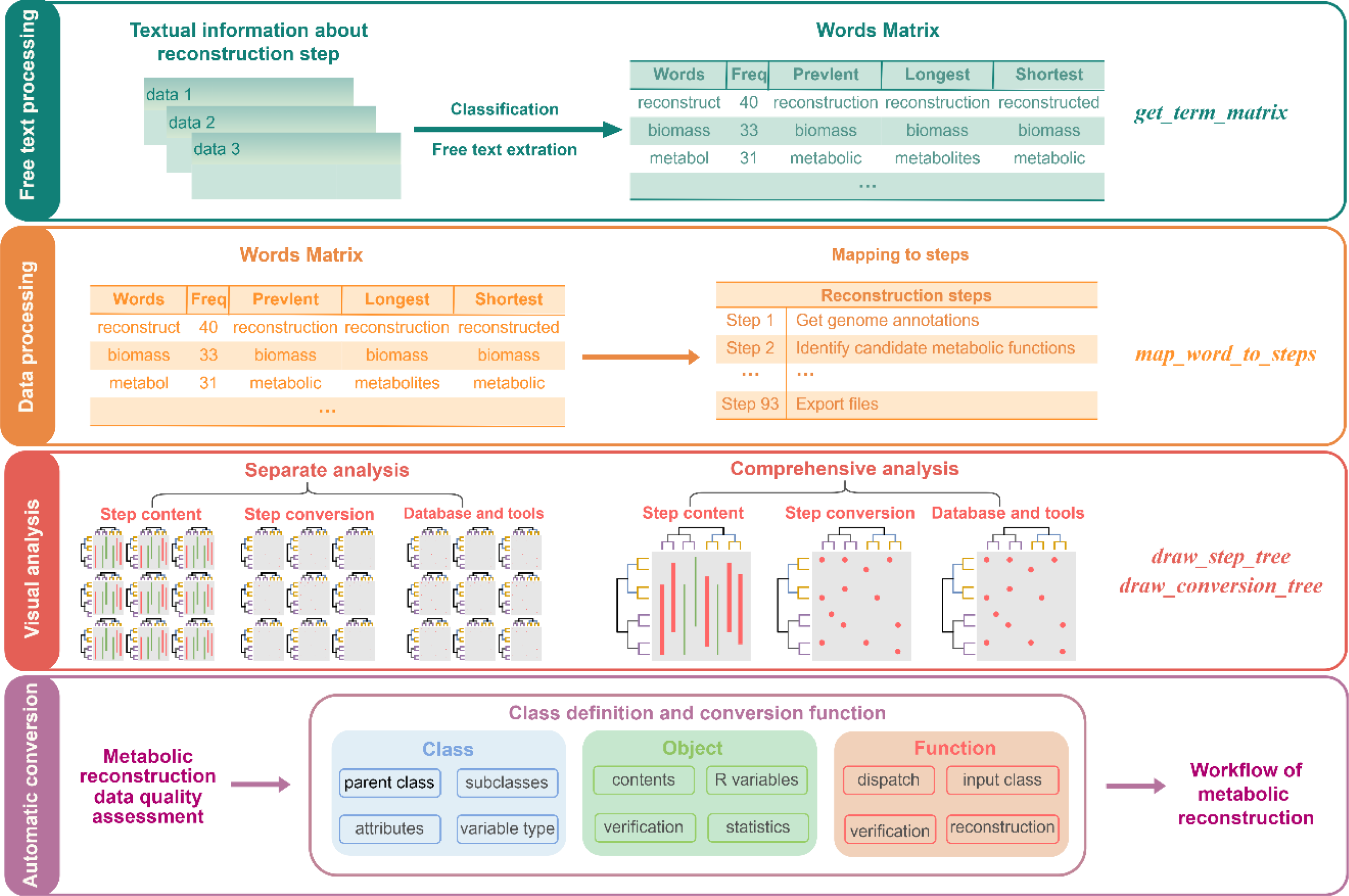
The workflow of Sysrecon. The *get_term_matrix* function converts the literature into the words matrix, and the *map_word_to_step* function is then utilized to map the words matrix to the corresponding steps of the metabolic reconstruction. Finally, the *draw_step_tree* and *draw_conversion_tree* functions are employed to generate visual representations of the mapped GSMR steps. Subsequently, the workflow of the GSMR is generated using the conversion function.

### Curating marker words

In order to minimize false positives and condense textual information for subsequent analytical processes, we manually assigned marker words to each step (**Supplementary data**). In text analysis, marker words are commonly employed for tasks such as categorization, clustering, sentiment analysis, and information extraction. For instance, the words, ‘obtain’ and ‘metabolites’, were used as the marker words of the step, obtain candidate metabolites. When selecting the marker words, the content and quantity of the marker words must be carefully determined. Broadly defined or too few marker words may lead to false positives when mapping to the GSMR step.

### Calculating mapping process

To assess whether the step was successfully map, we first established manually a mapping threshold for each GSMR step (**Supplementary data**). The mapping threshold is typically a binary decision variable, meaning that when the mapping result’s score above the threshold, a successful match has occurred; otherwise, a failed match has occurred. The mapping threshold is established in relation to the marker words during the mapping process. For instance, the marker words candidate and function must appear simultaneously in the step that identify candidate metabolic functions, in which case the mapping threshold is set to 1. Using the *map_word_to_step* function, we determined the metabolic reconstruction mapping score for each step:

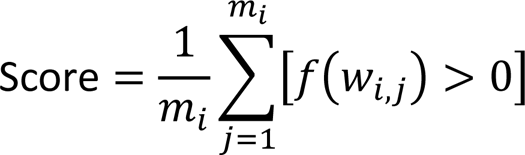

The variable *m_i_* represents the total number of marker words in the *i*^*th*^ step, while *w*_*i,j*_ denotes the j^th^ marker word in the *i*^*th*^ step. The function f(*w_i_*_,*j*_) represents the number of occurrences of the word *w*_*i,j*_ in the literature, and [*f*(*w*_*i,j*_) > 0] is the indicator function that takes the value of 1 when[*f*(*w*_*i,j*_) > 0], and 0 otherwise.

Given that some GSMR steps were frequently addressed in the literature, we took into account the significance of the convenience phases. When measuring the significance of steps, a marker word may correspond to several word forms in a literature. For example, the marker word pathway can correspond to pathwaytools, subpathway. To avoid information duplication, the marker word with the fewest occurrences was considered for the calculation when a marker word has several word forms.

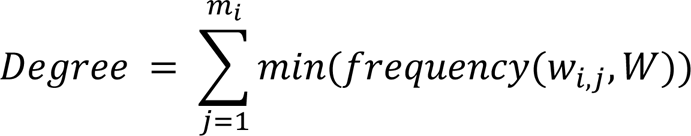

*W* represents all the words in the words matrix and *min*(*frequency*)) denotes the minimum frequency of each marker words *w*_*i,j*_ in *W*.

### Calculating the correlation of steps

In reconstruction, some steps are correlated, which is closely related to the metabolic function of the organism. In order to analyze the correlation between two metabolic processes, we need to quantify the data, transformation methods and tools required for each step. Thus, we refer to the stoichiometric matrix where -1 represents the input and 1 represents the output. Then, we calculated the correlation of two steps using the Euclidean distance.

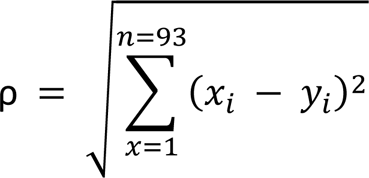

*x* and *y* represent any step in GSMR, *i* represents the data, transformation and tools needed. The results are then clustered via the *hculst* function with default parameters.

### Variables of systematic metabolic reconstruction

Systematic metabolic reconstruction was classified into eight phases, containing 93 steps in total (**Figure 3**). Thirty-six steps cannot be expressed in formulaic language, because most of them are manual operation. Each of the remaining 57 GSMR steps is defined by using a chemical formula composed of the Sysrecon variables which are the data, transformation methods, and tools needed in metabolic reconstruction. Each Sysrecon variable has unique abbreviation, to enhance the legibility and comprehensibility of the formulas.

**Figure 3:**
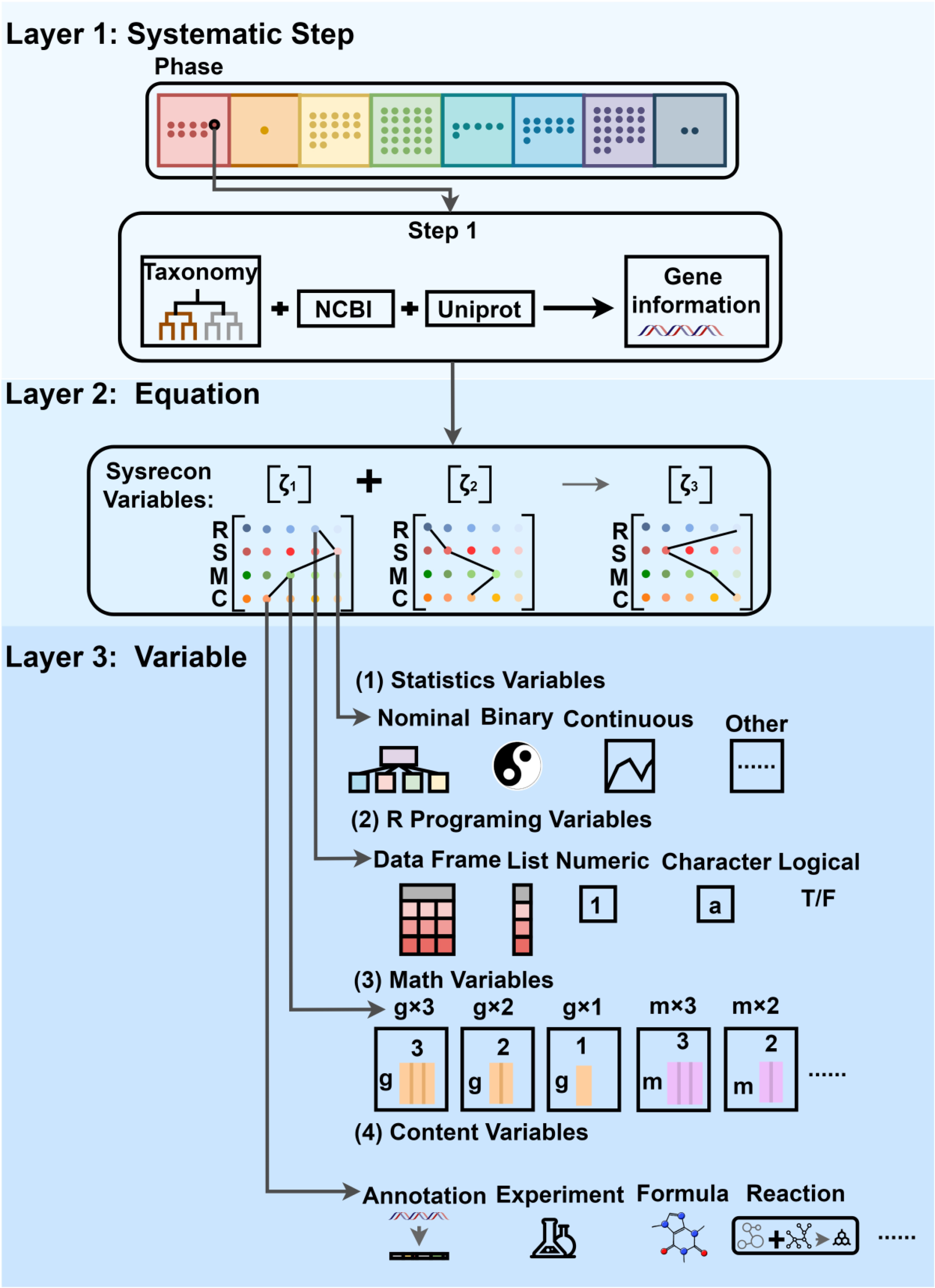
The flow chart of the variables of systematic metabolic reconstruction. The systematic metabolic reconstruction is consisted of 93 steps organized into 8 phases. These phases are denoted by different colors, and their respective meanings from left to right are data collection, draft reconstruction, data curation, reconstruction refinement, data quality control, information analysis, model evaluation, and reconstruction output. The step is represented in a format similar to a chemical formula, composed of various Sysrecon variables.

We partitioned the Sysrecon variables more precisely in order to expound on their content based on their data structure, content description, data dimensions, and statistical perspective. R variables are data structure-related variables that explain Sysrecon variables in terms of typical R data structures, which include data frames, lists, data, strings, and logical values. Content variables are related to the description content. It categorizes the Sysrecon variables from the point of view of describing Sysrecon, and categorizes the Sysrecon variables into elements such as reaction, annotation. Math variables depend on the number of dimensions in the data, and most Sysrecon variables are either 1 or 2 dimensional. Statistical variables are often divided into nominal, binary, continuous, and other variables and focus primarily on the Sysrecon content type.

### Conversion function

We utilized R programming for the implementation of a custom class, denoted as *systemrecon*, to organize and manage various elements of the reconstruction process. Firstly, we defined a function called slot, the primary objective of which is to store the Sysrecon variables and their elements mentioned in the literature into a list named *Info*. Then, Additional documentation was provided for the *systemrecon* class, specifying the Sysrecon variable name, concept, and a detailed list of slots (attributes) that the class contains. Each slot was described with its name, data type, and a brief description of its purpose within the class. Then, the *slotInfo* list was initialized and populated with information extracted from the Sysrecon varaiables using the slot function. Finally, the *systemrecon* class was formally defined by using the *setClass* function and specified the slots based on the *slotInfo* list.

When using *systemrecon* class, first we initiated an instance of the class, denoted as *test*, to serve as a container for storing and organizing data relevant to metabolic network reconstruction. The *test* will be populated with attributes and their associated information as the analysis progresses. Then, we maintained a list to store detailed descriptions of each reconstruction step and the associated attribute information. Additionally, we initialized an index variable, *i*, to keep track of the position of GSMR’s step in the list. Subsequently, we employed conditional statements to check for the presence of specific reconstruction steps within the results from *map_words_to_step* function. If a given step, such as step1, was identified in the data, we proceeded to initialize the corresponding attribute, such as Genetic_information, within the test, and the result and its position would be stored in the list. Finally, the list results were output to the screen.

## Results

### Integrated analysis of reconstruction data

Texts from 10 archaeal GSMR literature sources were compiled into a single text file for the analysis of their input data content. There are 31 steps related to input data that constitute the majority of archaeal metabolic reconstruction. The most prominent steps include identifying the main reactions, obtaining metabolites from biomass reactions, assembling draft reconstruction, adding demanding reactions for biomass reactions, and maximizing the objective functions of single secretion (**Figure 4**). Next, we integrated a total of 110 bacterial literature for analysis. We found that bacterial metabolic reconstruction involves a total of 54 key steps in reconstructing the data (**Supplementary Figure 1A**). The top five most frequently occurring steps in this process are identifying the main reactions, obtaining the metabolites from biomass reaction, assembling draft reconstruction, setting objective functions of biomass reactions, and maximizing the objective reactions of multiple secretions. Finally, a compilation of 55 literature sources on eukaryotic organisms was consolidated into a single text file for data input analysis. Eukaryotic metabolic reconstruction primarily encompasses 51 distinct steps, with the top five most frequently encountered steps: verifying the matrix, determining the reaction directionality, simulating the rich media, assembling draft reconstruction, and simulating the single-reaction deletion (**Supplementary Figure 1B**). These analyses showed that the steps of metabolic reconstruction still vary in different species, and that different steps may have different importance in subsequent GSMRs.

**Figure 4:**
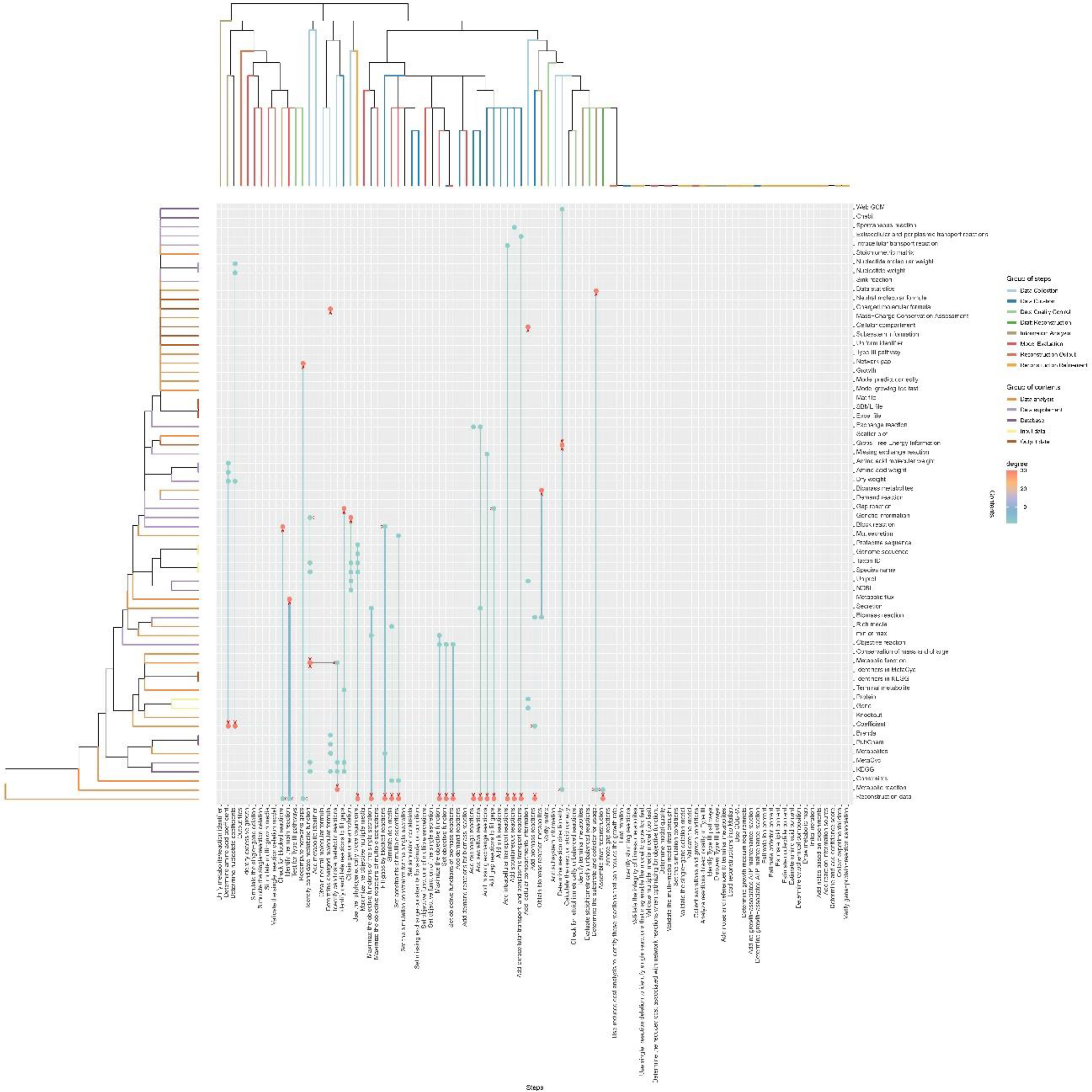
Results of the integrated analysis of the archaeal literature on the data of the reconstruction steps. The row shows the complete reconstruction steps involved in GSMR, and the column shows the input and output for the reconstruction steps. Green dots are input data, and red dots are output data. The thickness of the line and the shade of the color reflect the amount of information describing the step in the literature.

With our clustering analysis of metabolic step information and Sysrecon variables, we identified a cluster of 36 metabolic reconstruction steps that defied specific categorization, which were positioned on the far right of the dendrogram, and observed the grouping of similar metabolic steps, particularly those involving the addition of various metabolic reactions.

In the GSMR of these three organisms, there were 30 common steps (**Figure 5A**). This suggests that these steps are essential to the GSMR process of any organism. **Table 1** provides detailed information on these steps. Besides, there are specific reconstruction steps observed for different organisms. For instance, in bacterial GSMR, three steps are unique: calculating the reaction stoichiometry, adding metabolite identifiers, and checking for stoichiometrically unbalanced reactions. Eukaryotic GSMRs have only one GSMR step not available in the other two types of GSMRs, that is identifying whether the model grows too fast.

**Figure 5:**
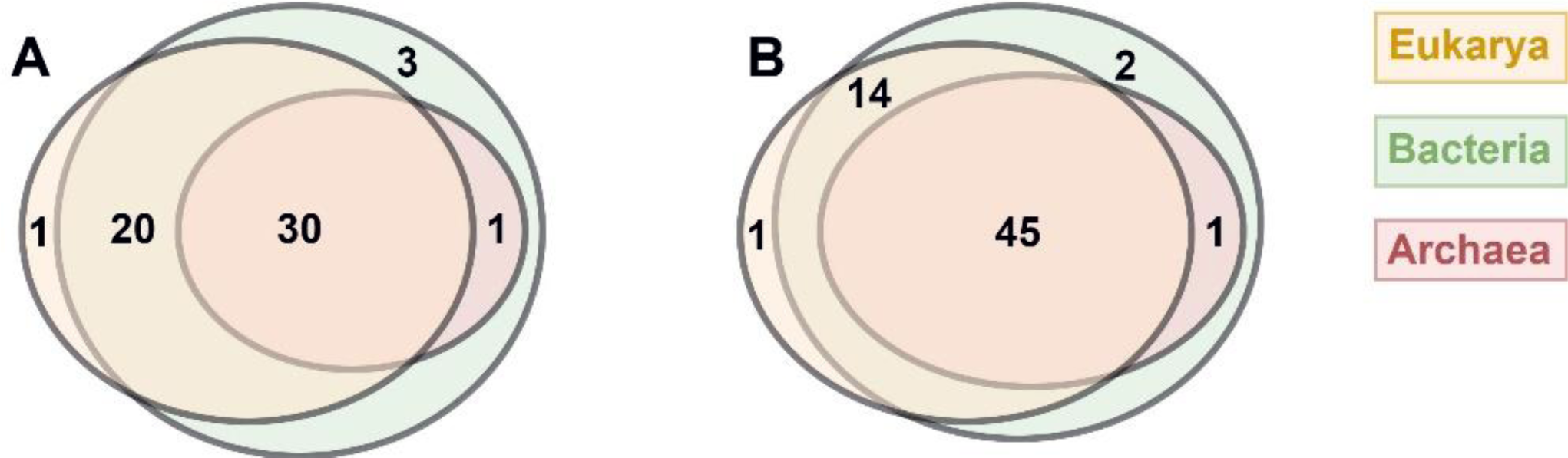
The integrated analysis of reconstruction steps and reconstruction step content for archaea, bacteria and eukaryotes. A: Venn diagram analysis of reconstruction steps that can be mapped to each type of species. B: Venn diagram analysis of Sysrecon variables that can be mapped to each type of species.

**Table 1:**
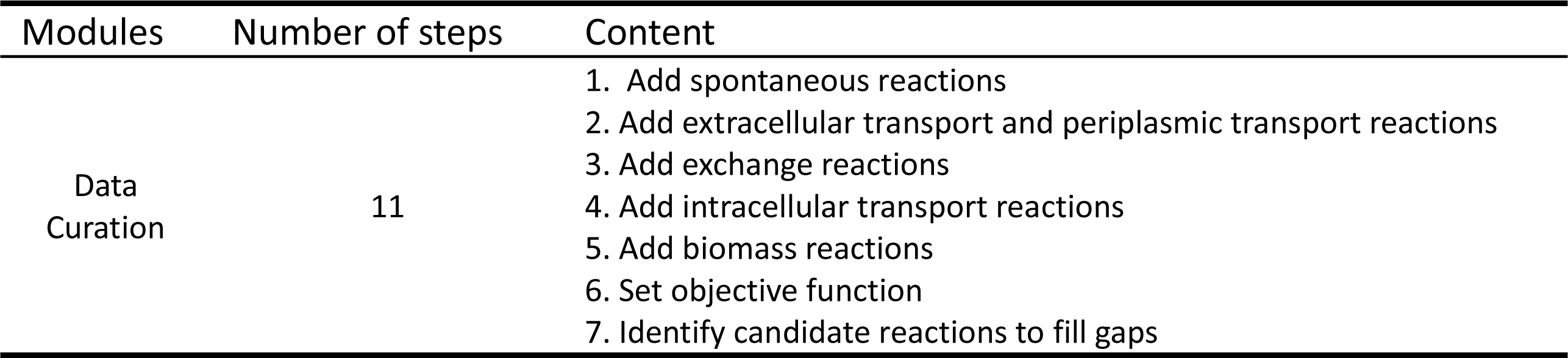
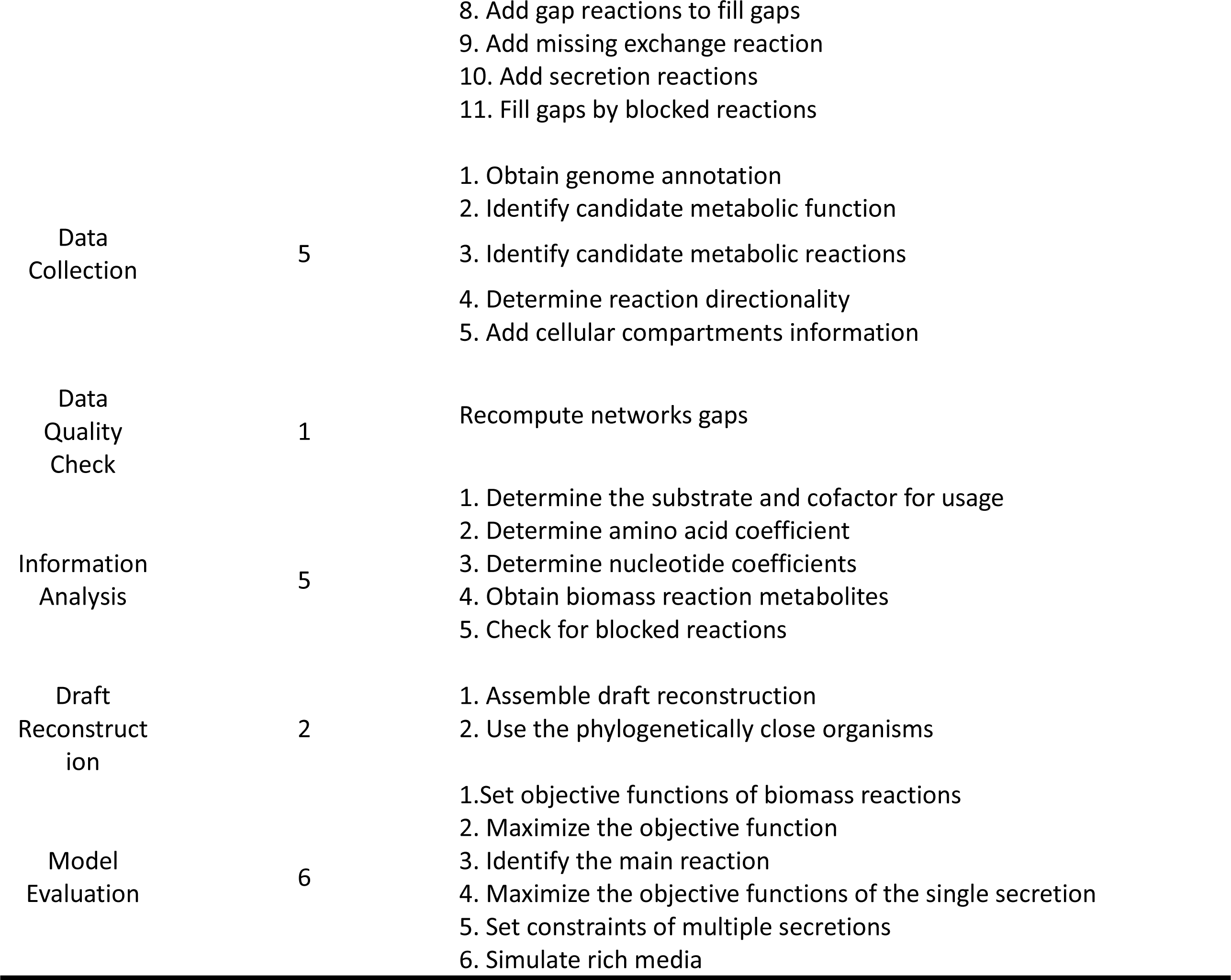
Reconstruction steps for GSMR.

| Modules | Number of steps | Content |
| --- | --- | --- |
| Data Curation | 11 | 1. Add spontaneous reactions<br>2. Add extracellular transport and periplasmic transport reactions<br>3. Add exchange reactions<br>4. Add intracellular transport reactions<br>5. Add biomass reactions<br>6. Set objective function<br>7. Identify candidate reactions to fill gaps |
|  |  | 8. Add gap reactions to fill gaps<br>9. Add missing exchange reaction<br>10. Add secretion reactions<br>11. Fill gaps by blocked reactions |
| Data Collection | 5 | 1. Obtain genome annotation<br>2. Identify candidate metabolic function<br>3. Identify candidate metabolic reactions<br>4. Determine reaction directionality<br>5. Add cellular compartments information |
| Data Quality Check | 1 | Recompute networks gaps |
| Information Analysis | 5 | 1. Determine the substrate and cofactor for usage<br>2. Determine amino acid coefficient<br>3. Determine nucleotide coefficients<br>4. Obtain biomass reaction metabolites<br>5. Check for blocked reactions |
| Draft Reconstruction | 2 | 1. Assemble draft reconstruction<br>2. Use the phylogenetically close organisms |
| Model Evaluation | 6 | 1. Set objective functions of biomass reactions<br>2. Maximize the objective function<br>3. Identify the main reaction<br>4. Maximize the objective functions of the single secretion<br>5. Set constraints of multiple secretions<br>6. Simulate rich media |

We also conducted an analysis of Sysrecon variables in the metabolic reconstructions of archaea, bacteria, and eukaryotes (**Figure 5B**). There are 45 common Sysrecon variables and various Sysrecon variables specific for three types of organisms. Among them, two variables are distinctive to bacteria, such as the molecular formula from CHEBI database and stoichiometrically unbalanced reactions, while only one Sysrecon variable is exclusive to eukaryotes, the model growth rate. The results of metabolic step analysis support the results from Sysrecon variable analysis.

### Integrated analysis of transformation methods

In our analysis of archaea, we found that 35 metabolic reconstruction steps involve transformation methods (**Figure 6**). We conducted a frequency analysis of these steps, and the top five most frequent steps include identifying the main reactions, assembling the draft reconstruction, setting objective functions of biomass reactions, and maximizing the objective functions of single secretion. After analyzing the step transition information in the literature related to bacterial metabolic reconstruction, we identified a total of 58 metabolic reconstruction steps related transformation methods (**supplementary figure 2A**). The top five most frequent ones in the literature are the identification of terminal metabolites, the inclusion of demand reactions for biomass reactions, the identification of main reactions, the maximization of objective functions for single secretion, and the maximization of objective reactions for multiple secretions. Finally, in the eukaryotic organisms, we discovered a total of 55 metabolic reconstruction steps that require transformation methods (**supplementary figure 2B**), with the top five most frequent reconstruction steps including the validation of stoichiometric matrix, determination of reaction directionality, simulation of rich media conditions, assembly of draft reconstructions, and the inclusion of information sources for reactions in the analysis for step transformation information. By analyzing the steps of the transformation methods, it becomes evident that these steps bear a significant resemblance to the requirements of previous GSMRs. This indicates that in GSMRs, attention must be paid to the data transformation methods for each step.

**Figure 6:**
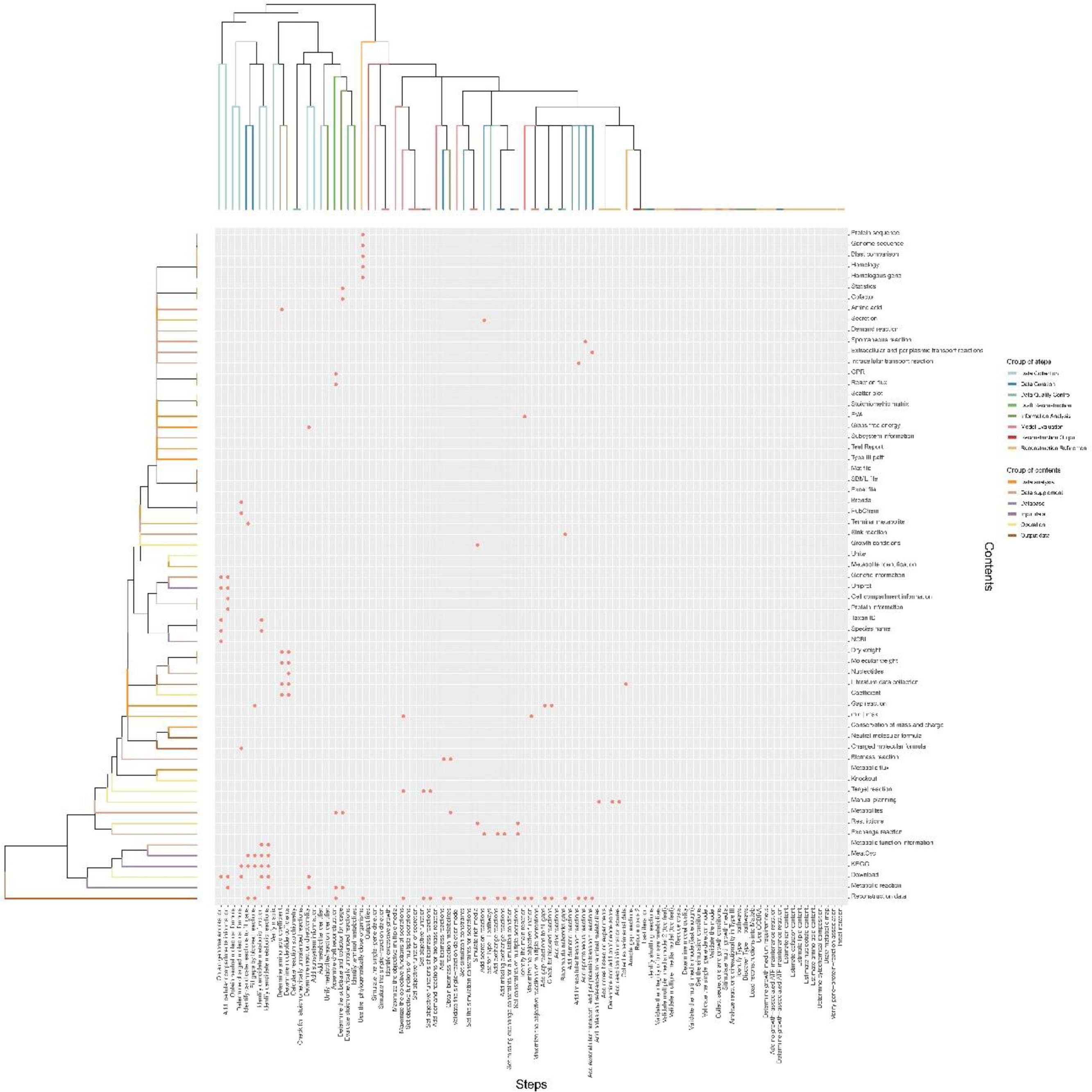
Results of the integrated analysis of the archaeal literature on step transformation information. Columns show the complete reconstruction steps involved in the GSMR, and rows show the transformation information involved in the reconstruction steps. The dots represent the details of the transformation information involved in each reconstruction step.

A clustering analysis of reconstruction transformation methods and related Sysrecon variables was also conducted. During this process, we observed some trends. Firstly, those reconstruction steps that couldn’t be precisely defined were clustered on the right side of the tree, indicating their unique characteristics within the overall metabolic reconstruction. Meanwhile, modules associated with data collection, such as obtaining genome annotation information, acquiring cellular compartment details, and gathering neutral molecular formula data, were consolidated on the left side of the tree, underscoring their significance in the information acquisition process. This result is also broadly consistent with the previous cluster analysis.

We conducted a comprehensive analysis of the procedures related to transformation methods in the literature of archaea, bacteria, and eukaryotes. The findings revealed a total of 34 metabolic steps among them (**Figure 7A**). This represents an additional four steps compared to the comprehensive analysis of reconstruction data. These additional four reconstruction steps encompass the collection of experimental data, the addition of confidence scores, the annotation of metabolic reaction information sources, and the annotation of terminal metabolites. It is noteworthy that these particular steps, due to the unavailability of their Sysrecon variables, have been categorized as manually devised procedures within the analysis.

**Figure 7:**
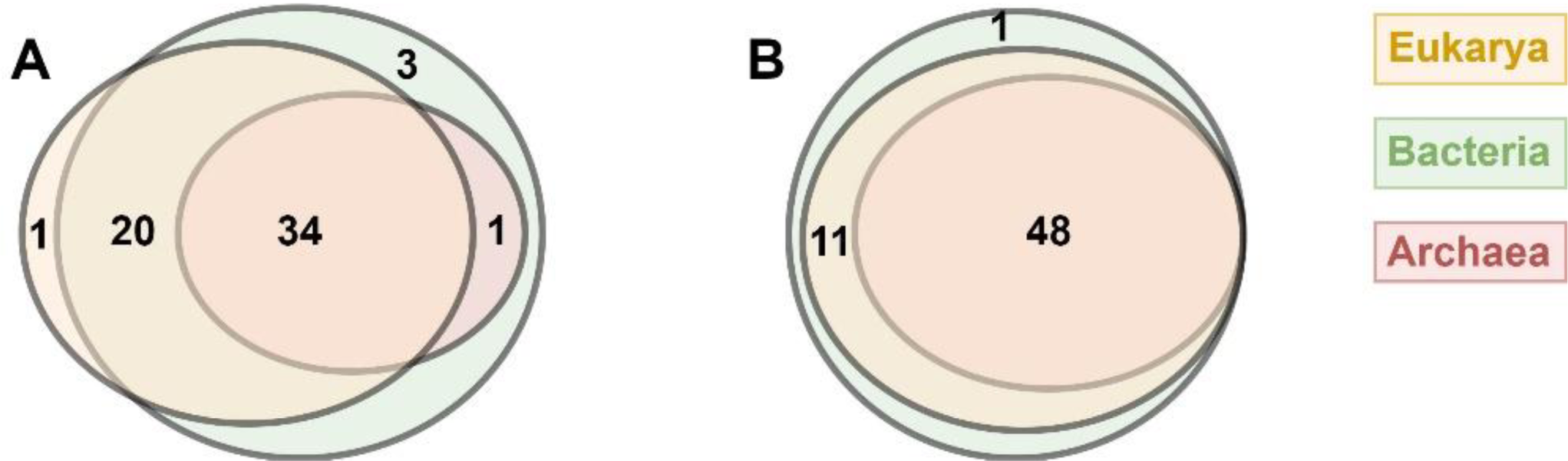
The integrated analysis of reconstruction steps and Sysrecon variables for archaea, bacteria and eukaryotes. A: Venn diagram analysis of reconstruction steps that can be mapped to each type of species. B: Venn diagram analysis of Sysrecon variables that can be mapped to each type of species.

The Sysrecon variables result showed that bacterial, archaeal, and eukaryotic metabolic reconstruction methods share similar transformation approaches (**Figure 7B**). Nonetheless, bacteria exhibit one specific Sysrecon varibales result not present in the other two types of organisms: the generation of model assessment reports.

### Integration analysis of tools

The literature analysis of archaeal metabolism reveals a total of 46 steps associated with reconstruction tools (**Figure 8**). Among the most frequently occurring GSMR steps, identifying main reactions, obtaining metabolites from biomass reactions, assembling draft reconstructions, setting objective functions of biomass reactions, and maximizing the objective functions of secretions are prevalent. Similarly, our analysis of bacterial GSMR literatures explore the databases and tools required for bacterial metabolism reconstruction. The results indicated that 84 reconstruction steps require tools (**Supplementary Figure 3A**). The five most commonly employed steps across all literature include identification of terminal metabolites, adding demand reactions for biomass reaction, identification of the main metabolic reactions, as well as the optimization of secretion objective functions, both single and multiple, aligning with the preceding two analyses. After analyzing literature on eukaryotic metabolic reconstruction, we identified a total of 80 reconstruction steps associated with the tools (**Supplementary Figure 3B**). Among these, the top five most frequently used steps were verifying stoichiometry, determining reaction directionality, simulating the rich media, assembling draft reconstruction, and adding reaction information sources. When analyzing the steps related to tools and metabolic methods, we found that they are roughly consistent with the steps of inputting data, which shows that these three factors are interrelated, and in the subsequent metabolic reconstruction process. Also, we need to consider the data of each step, the method of transformation, and the tools needed, which also shows that there is a consistency.

**Figure 8:**
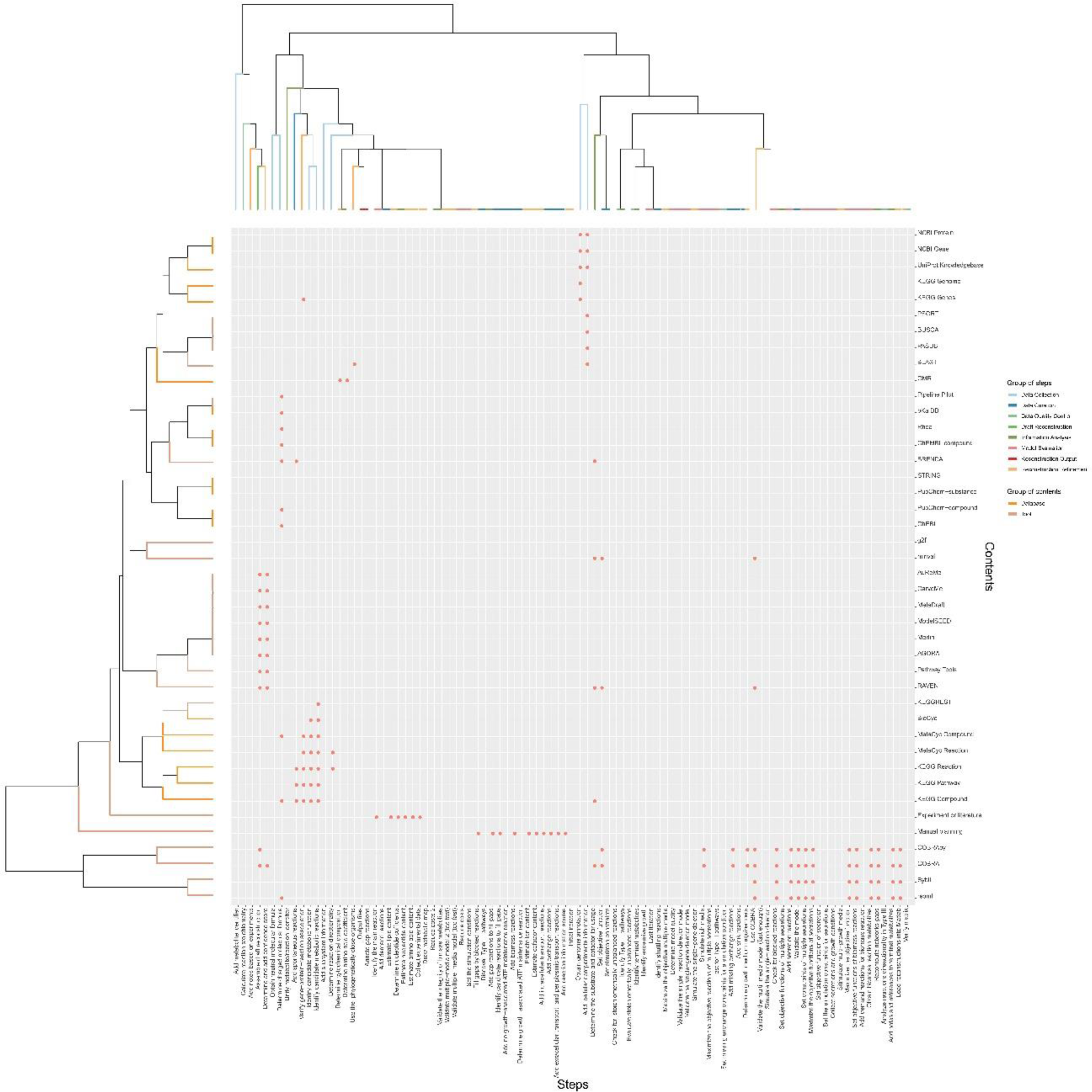
The integrated analysis of tools information for archaeal literature. Columns show the complete reconstruction steps involved in the GSMR, and rows show the database and tool information involved in the reconstruction steps. The dots represent the details of the database and tool information available for each reconstruction step.

**Figure 9:**
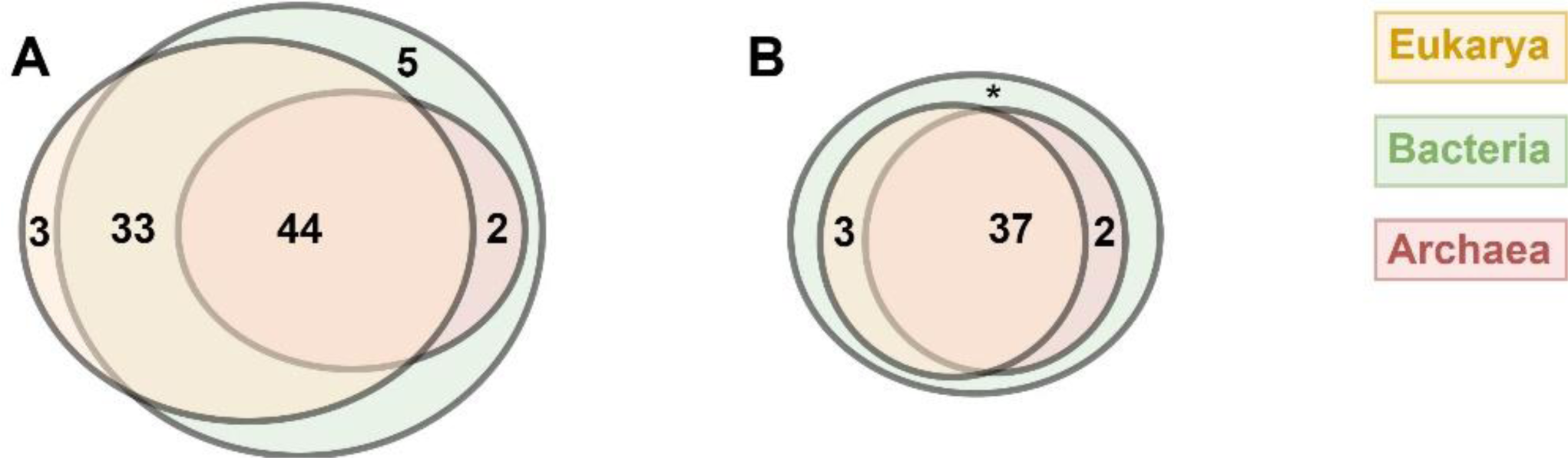
The integrated analysis of reconstruction steps and tools information for archaea, bacteria and eukarya. A, Venn diagram analysis of the reconstruction steps to which each type of species can be mapped. B, Venn diagram analysis of the tools information to which each type of species can be mapped. * indicates that the tools used for eukarya are not specific.

However, a cluster analysis of the reconstruction steps with tools information demonstrated that no specific modules are grouped together, even for those reconstruction steps that are unsure which databases and tools were employed. This suggests that even for similar reconstruction steps, distinct tool information is employed.

Through a comprehensive analysis of the tools from literature on archaea, bacteria, and eukaryotes, it was found that 44 reconstruction steps are common. Among these, bacteria have five unique steps, including validating the conservation of mass and charge, adding metabolite identifier, initial iteration to refine the reconstruction data, evaluating stoichiometrically unbalanced reactions, and last iteration to refine the reconstruction data. Eukaryotic-specific steps encompassed collecting secretion and growth conditions, identifying shuttle reactions, and determining whether the model is growing too fast.

Regarding tools usage, there are 37 tools that could be employed in GSMRs across these three biological categories. However, there are less specific tools exclusive to any of them, indicating that these biological metabolic reconstructions typically utilize generic tools.

### Comparison of plant GSMR models

We collected 18 articles about plant GSMRs, and then gathered information on the metabolites, metabolic reactions, and compartments (Botero et al., 2018; Chatterjee et al., 2017; Cheung et al., 2013; de Oliveira Dal’Molin et al., 2010; Gerlin et al., 2021; Grafahrend-Belau et al., 2009; Hay and Schwender, 2011; Kim et al., 2016; Moreira et al., 2019; Pfau et al., 2018a; Pilalis et al., 2011; Poolman et al., 2013; Saha et al., 2011; Seaver et al., 2015; Shaw and Cheung, 2018; Simons et al., 2014; Yuan et al., 2016). We observed that as the years passed, the data volume in some reconstructions increased, and the information became more comprehensive. However, due to the specific requirements of certain models, their information may not be entirely complete.

The data presented in **Table 2** elucidates variations arising from disparities in model content. To gain a more in-depth understanding of these divergences, we plan to conduct a Sysrecon analysis on these models.

**Table 2:**
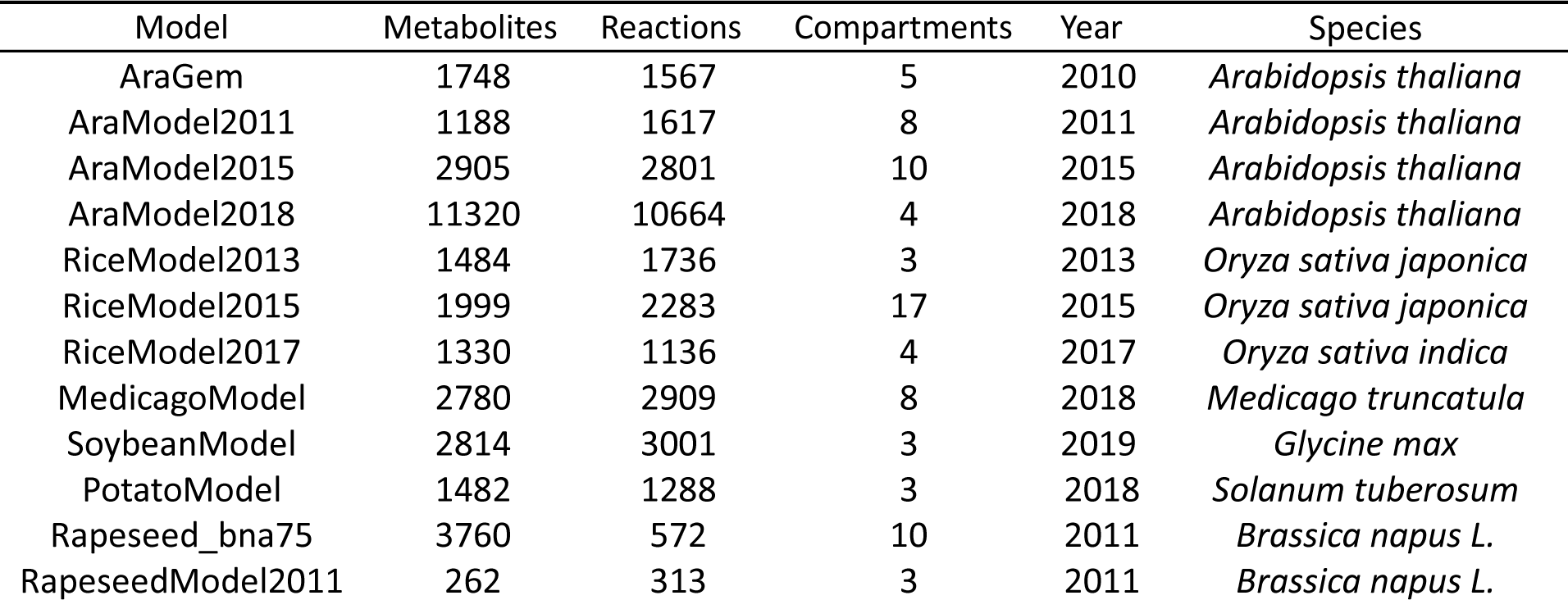
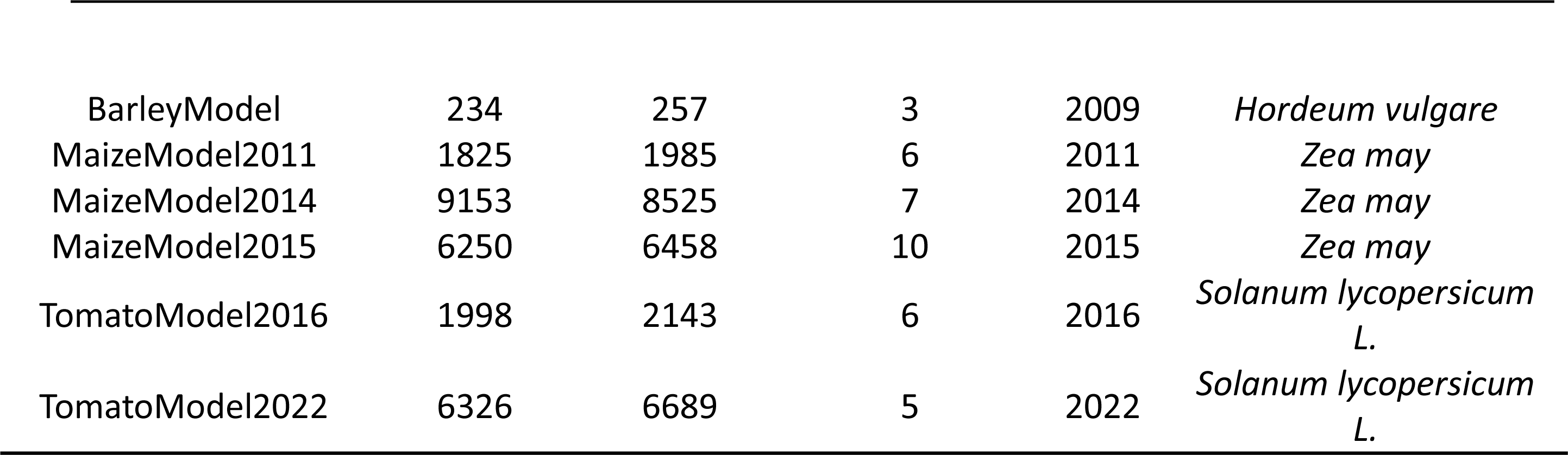
Information on the GSMR of different plant types.

| Model | Metabolites | Reactions | Compartments | Year | Species |
| --- | --- | --- | --- | --- | --- |
| AraGem | 1748 | 1567 | 5 | 2010 | *Arabidopsis thaliana* |
| AraModel2011 | 1188 | 1617 | 8 | 2011 | *Arabidopsis thaliana* |
| AraModel2015 | 2905 | 2801 | 10 | 2015 | *Arabidopsis thaliana* |
| AraModel2018 | 11320 | 10664 | 4 | 2018 | *Arabidopsis thaliana* |
| RiceModel2013 | 1484 | 1736 | 3 | 2013 | *Oryza sativa japonica* |
| RiceModel2015 | 1999 | 2283 | 17 | 2015 | *Oryza sativa japonica* |
| RiceModel2017 | 1330 | 1136 | 4 | 2017 | *Oryza sativa indica* |
| MedicagoModel | 2780 | 2909 | 8 | 2018 | *Medicago truncatula* |
| SoybeanModel | 2814 | 3001 | 3 | 2019 | *Glycine max* |
| PotatoModel | 1482 | 1288 | 3 | 2018 | *Solanum tuberosum* |
| Rapeseed_bna75 | 3760 | 572 | 10 | 2011 | *Brassica napus L.* |
| RapeseedModel2011 | 262 | 313 | 3 | 2011 | *Brassica napus L.* |
| BarleyModel | 234 | 257 | 3 | 2009 | *Hordeum vulgare* |
| MaizeModel2011 | 1825 | 1985 | 6 | 2011 | *Zea may* |
| MaizeModel2014 | 9153 | 8525 | 7 | 2014 | *Zea may* |
| MaizeModel2015 | 6250 | 6458 | 10 | 2015 | *Zea may* |
| TomatoModel2016 | 1998 | 2143 | 6 | 2016 | *Solanum lycopersicum*<br>L. |
| TomatoModel2022 | 6326 | 6689 | 5 | 2022 | *Solanum lycopersicum*<br>L. |

The findings in **Figure 10A** uncovered that the majority of GSMR steps of the same plant gradually increase, indicating an improvement in quality. Nevertheless, some reconstructions exhibit a decrease in the number of steps (e.g., AraModel2015 and AraModel2018). The decrease could be attributed to specific purposes of GSMR, where certain aspects of plant metabolism, such as photosynthesis and respiration, are given priority, leading to a reduction in the number of steps. Furthermore, species differences could also contribute to variations in the steps and quality of GSMRs. For instance, subspecies of rice, such as *indica* and *japonica*, may display distinct steps and quality levels in their respective GSMRs.

**Figure 10:**
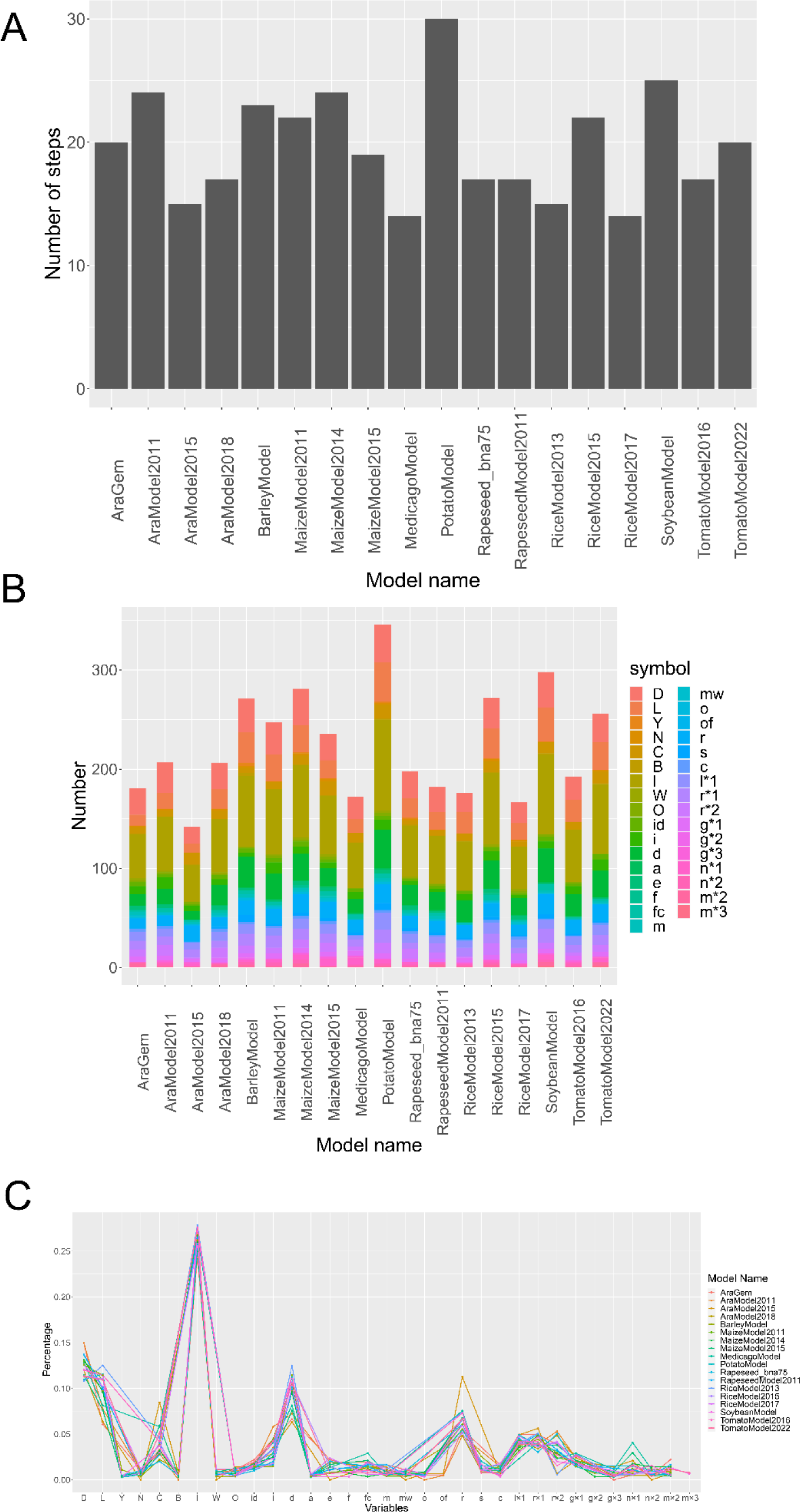
Visualization of GSMR steps and variables. A. Bar chart of metabolic reconstruction versus metabolic steps; B. Bar chart of total number of elements in metabolic reconstruction; C. Line graph of metabolic reconstruction versus elemental occupancy.

In accordance with results from the reconstruction steps of different species’ metabolism, we have initiated our focus on the elements involved in these metabolic reconstruction steps. Given that each Sysrecon variable comprises R programming variables, statistical variables, math variables, and content variables, and these variables encompass various elements, we have redirected our attention from variables to elements (**Figure 1**). Firstly, we computed the total number of elements in each metabolic reconstruction (**Figure 10B**). The results indicated a direct proportionality between the total number of variables and the number of GSMR steps, meaning that as the number of metabolic reconstruction steps increases, the total number of variables also increases. However, it is imperative to consider the diversity of variable types within each reconstruction. A greater variety of variable types suggests a broader range of information encompassed within that reconstruction.

We conducted a quantitative analysis of each reconstructed element type, and the results indicate that the majority of element types number around 25. (**Supplementary Figure 4**). Nevertheless, some reconstructions exhibited a dearth of element categories, which can be ascribed to several factors. However, there are still some reconstructed element types with fewer instances, which can be attributed to several factors. Firstly, some earlier publications may have limited steps due to insufficient information, thereby restricting the diversity of element types. Secondly, data for certain species may be relatively scarce, leading to a reduced requirement for element types in metabolic reconstructions. Lastly, it is important to note that GSMRs typically have specific focuses, concentrating solely on particular metabolic processes, which may result in a lack of other relevant data during the reconstruction process.

Next, we computed the proportions of each variable within each model and depicted them using line plots (**Figure 10C**). The results reveal a consistent overall trend in variable proportions across all models.

In R programming variables, data frame elements have the highest proportion, primarily due to their significant role in metabolic reconstruction. These variables encompass crucial information, including metabolic reactions and chemical formulas of substances. This information is not only extensive but also pivotal, and its absence could potentially diminish the quality of the reconstruction.

Among the statistical variables, nominal variable elements have the highest proportion. This is primarily because nominal variable elements in metabolic reconstruction are typically used to represent various attributes of organisms, such as different species, reactions, compartments. Given the extensive information involved in metabolic reconstruction, nominal variables serve to categorize and compare metabolic networks. Therefore, there is a consistent trend of nominal variables’ proportion across all models.

In terms of content variables, data elements and reaction elements both have approximately equal proportions at the top. The high proportion of data variables is primarily attributed to the active participation of reconstruction data in the GSMR process. This is primarily evident in different metabolic reconstructions, where reconstruction data is continuously updated with new content and iterated to generate a high-quality metabolic reconstruction. As for reaction elements, their high proportion can be attributed to two main factors. Firstly, reaction variables are categorized into various types, such as metabolic reaction, spontaneous reactions, and exchange reactions. Secondly, these reaction elements have a higher frequency of occurrence in the metabolic reconstruction steps, as most metabolic reconstructions require a variety of reactions for supplementation.”

The element with the highest frequency among the math variables is *r × 1*, primarily due to the prevalence of this unit in elements comprising data frame elements, nominal variables elements, and reaction elements. This observation is consistent with the trend in the percentage of a forementioned three variables.

Subsequently, we examined the relationship between elements and metabolic reconstructions, particularly for the same species across different years. We employed a *chi*-squared test to assess whether the four Arabidopsis metabolic reconstructions from 2009 to 2018 exhibited significant differences in element composition. The *chi*-square test yielded a *p*-value greater than 0.995, indicating that the elemental content differences between the reconstructions were not significant. This observation implies a high degree of similarity among these four metabolic reconstructions, possibly sharing similar elemental components within metabolic pathways. This finding also confirms the consistent trend in the proportions of most elements across different reconstructions, indicating an overall consistency in plant metabolic reconstructions. Therefore, when conducting metabolic reconstructions for Arabidopsis or other plant species, evaluating the normality of reconstruction steps through element quantity and proportions can aid in further optimizing the metabolic reconstruction process or exploring whether the species possesses unique biological characteristics.

## Discussions

Numerous GSMR software or toolkits are widely utilized in scientific research, but they still have some shortcomings. For instance, the COBRA toolbox is grounded on the constrained model of metabolic flux balance and steady-state assumptions, which are primarily used for the static analysis of metabolic networks. This approach cannot analyze the dynamic response and temporal regulation of metabolic networks. RAVEN toolbox is founded on the dynamic metabolic network model, which presents higher computational complexity, necessitating longer computation time and high-performance computing resources. Additionally, GPRuler has limited functionality in collecting GPR information of species (Di Filippo et al., 2021). Conversely, MERLIN employs a visual interface that relies on precise biological information, including metabolic pathways, enzyme-catalyzed reactions, and metabolites, among others. Nonetheless, inaccurate or imprecise biological information can negatively impact the accuracy of MERLIN’s analysis results. It is important to note that the reliability of the final results obtained by MERLIN is heavily influenced by the accuracy of the input data.

Due to the inconsistent quality of GSMRs generated by different tools, Sysrecon is utilized to perform a comprehensive evaluation of the reconstruction steps across different reconstructions. Firstly, Sysrecon provides a standardized metabolic reconstruction framework that allows different GSMRs to follow a unified workflow. Additionally, Sysrecon evaluates the importance of reconstruction steps by calculating their frequency of occurrence across all literature. The reconstruction steps with higher frequencies are considered more important and may require repeated implementation for improving the reconstruction data. Sysrecon also provides detailed information on the implementation process of each reconstruction step, including operational and algorithmic information as well as data and tools utilized. Finally, Sysrecon allows for visualizing the analysis results, providing an intuitive comparison of different GSMRs in terms of the reconstruction steps and tools used.

By analyzing all collected GSMR literatures, we evaluated each reconstruction step and its related elements and constructed a high-quality systematic reconstruction assessment framework that can provide users with the necessary steps and related data for GSMR, as well as further refinement. First, we intend to employ manual documentation to increase the reliability and credibility of the results due to faults and omissions in computerized text analysis. Second, it is also crucial to establish a quality assessment system for databases and tools. Given the differences in the databases and tools used by each reconstruction process and the different coverage and data processing capabilities, the quality of the data produced will vary even for the same reconstruction process. Therefore, we plan to include a quality assessment system for databases and tools in the system reconstruction assessment framework.

In this study, we performed a comprehensive analysis of all available literature on GSMR and evaluated each step and related element of the reconstruction process to develop a high-quality, systematic framework for GSMR assessment. The framework provides users with essential steps and related data for GSMR, along with recommendations for further refinement. To improve the reliability and credibility of our results, we plan to incorporate manual curation of literature to address potential errors and omissions resulting from automated text analysis. In addition, we recognize the importance of establishing a quality assessment system for databases and tools used in the reconstruction process. Due to the differences in coverage and data processing capabilities of different databases and tools, the quality of the reconstructed data may vary significantly. Hence, we intend to include a quality assessment system for databases and tools in our reconstruction assessment framework.

## Conclusion

Currently, GSMR is rapidly advancing, providing a new perspective for research in the field of biology. This study created an assessment framework for different reconstructions. We summarized the steps of metabolic reconstruction and collected common data, transformation methods, and tools used in the metabolic reconstruction process. Subsequently, the Sysrecon package identified critical steps in metabolic reconstruction based on the literature, providing users with reference recommendations for subsequent reconstruction processes and specifying the required data and tools for each step, enhancing the reproducibility of models. Additionally, with the assistance of the Sysrecon package, we can determine whether a species’ metabolic reconstruction model exhibits significant specificity, allowing for more targeted data collection, data processing, and tool usage in the future. This package can assist GSMR and improve the quality of GSMR, contributing to the future development of GSMR.

## Key Points

1. Sysrecon provides a quantitative framework for defining GSMR steps, and can facilitate the development of high-quality GSMRs.
2. A comprehensive textual analysis of all published GSMR literature is performed to extract quantitative and qualitative step data.
3. A formula-based method is developed to decompose a GSMR procedure into many step blocks that can then be combined and modified according to the species-specific parameters.

## Data availability

To ensure the transparency and repeatability of systematic GSMR, we uploaded the relevant code and data to Bitbucket (https://bitbucket.org/ouyangshilin/sysrecon_code_r/src/master/) and Sysrecon is published on CRAN (https://cran.r-project.org/web/packages/Sysrecon/).

